# Carbon Capture Modeling and Simulation Platform: A Coupled Microalgal Bioreactor–Yeast Fermentation Approach for Bioethanol Production

**DOI:** 10.64898/2026.03.31.715672

**Authors:** Alaa Hamid, Nasma Akasha, Plamedie Kapenda Mukumbi, Ammar Mirghani, Tamir Omer

## Abstract

This article presents the development of an advanced modeling and simulation platform for carbon capture systems, with a focus on integrated process analysis from upstream CO_2_ capture through to bioethanol production. The platform supports the evaluation of CO_2_ mitigation technology by coupling mathematical bioprocess models with an interactive desktop application. The biological system employs Chlorella vulgaris microalgae to fix CO_2_ through photosynthesis and generate carbohydrate substrates, which are subsequently converted to bioethanol by Saccharomyces cerevisiae yeast via fermentation. The simulation integrates three established kinetic models—the Monod, Logistic, and Luedeking-Piret models—to predict biomass growth, substrate consumption, and ethanol yield under varying operational conditions. A closed-loop CO_2_ recycling subsystem captures fermentation off-gases and reintroduces them into the bioreactor, enhancing overall carbon utilization efficiency. Three representative simulation scenarios demonstrated process efficiencies ranging from 1.09% to 93.78% of the theoretical maximum CO_2_-to-ethanol conversion efficiency, confirming the platform’s capacity to evaluate a wide operational envelope. The Electron/React-based desktop application provides real-time visualization, interactive 3D bioreactor models, and a simulation history module, making it accessible to researchers, engineers, and students. The platform serves as a digital twin that bridges rigorous bioprocess mathematics with intuitive user interaction, providing a cost-effective tool for designing and optimizing sustainable carbon capture and biofuel production systems.

## 1. Introduction

The accelerating accumulation of anthropogenic carbon dioxide (CO_2_) in the Earth’s atmosphere represents one of the most critical environmental challenges of the twenty-first century. Atmospheric CO_2_ concentrations have risen from approximately 280 parts per million (ppm) in pre-industrial times to over 420 ppm today, with industrial processes, energy generation, and transportation identified as the primary emission sources [1]. This escalation is directly linked to global temperature increases, ocean acidification, and an increase in the frequency of extreme weather events, all of which underscore the urgent need for scalable, economically viable, and technically robust carbon management strategies.

Carbon Capture and Utilization (CCU) encompasses a broad portfolio of technologies aimed at intercepting CO_2_ before atmospheric release and converting it into valuable products. Among the most promising biological routes is the coupling of microalgal CO_2_ fixation with fermentative biofuel production. Microalgae are particularly well-suited for this purpose: they achieve photosynthetic efficiencies significantly higher than terrestrial plants, can be cultivated on non-arable land using industrial flue gas as a carbon feedstock, and can accumulate substantial quantities of carbohydrates and lipids suitable for downstream conversion into bioethanol, biodiesel, and biogas [1,2].

The microalga Chlorella vulgaris has been extensively studied in the context of carbon capture and biofuel production due to its high CO_2_ fixation rate (up to 1.5–2.5 g CO_2_/g dry biomass per day), its tolerance of elevated CO_2_ concentrations, and its ability to accumulate carbohydrate fractions suitable for fermentation [1,2]. When coupled with the yeast Saccharomyces cerevisiae—the workhorse organism of industrial bioethanol production—a two-stage biological system emerges: the photochemical bioreactor converts CO_2_ into glucose-rich biomass, and the downstream fermenter converts that glucose into ethanol and CO_2_. Crucially, the CO_2_ released during fermentation can be recycled back to the bioreactor, creating a partially closed-loop carbon cycle that improves overall system efficiency.

Despite the scientific and industrial interest in such systems, a significant gap exists between theoretical process design and accessible computational tools for system-level evaluation. Existing commercial simulators (e.g., SuperPro Designer, Aspen Plus) require specialized expertise and expensive licensing, while purely mathematical implementations lack user-friendly interfaces for rapid scenario exploration. The present work addresses this gap by developing a Carbon Capture Modeling and Simulation Platform—a desktop application that embeds validated kinetic models within an interactive visualization environment.

The platform integrates the Monod model for substrate-limited microbial growth [5], the logistic model for carrying-capacity-constrained biomass dynamics [6], and the Luedeking-Piret model for growth- and non-growth-associated product formation [7,8]. Environmental control algorithms simulate the effects of temperature, pH, light intensity, and nutrient availability, providing a multi-parameter assessment framework. The application was developed using an Electron/React/JavaScript frontend and a Python backend compiled to a portable executable, ensuring cross-platform compatibility without requiring users to install scientific computing environments.

This article describes the bioprocess design rationale (Section 2), the mathematical formulations underpinning the simulation engine (Section 3), the software architecture and technical implementation (Section 4), simulation results and validation scenarios (Section 5), and conclusions with recommendations for future development (Section 6). By combining technical rigor with accessibility, the platform aims to democratize the design and pre-feasibility assessment of integrated carbon capture and biofuel production systems.

**Figure 1.**
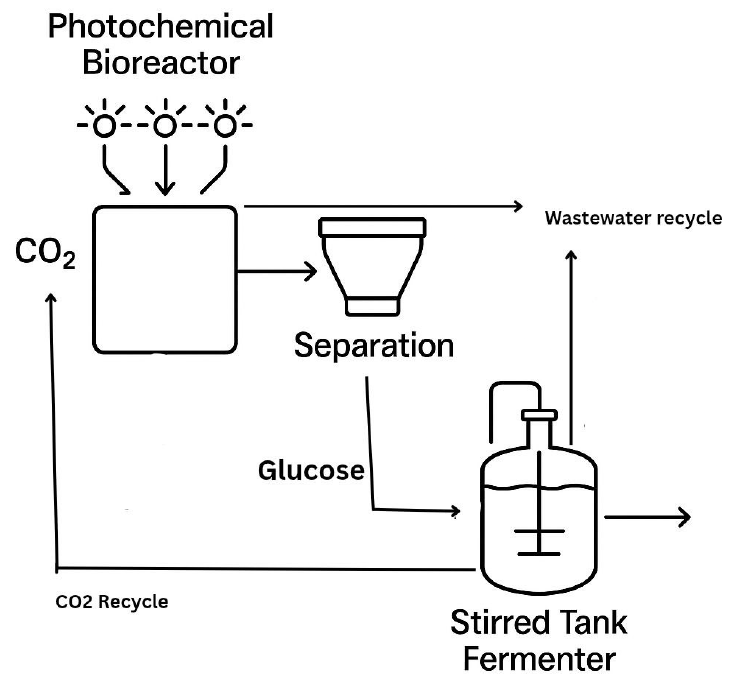
Conceptual schematic of the integrated CO_2_-to-bioethanol platform showing the photochemical bioreactor, fermenter, and CO_2_ recycling loop.

## 2. Materials and Methods

### 2.1. Organism Selection and Biological Rationale

The selection of biological agents for a CO_2_-to-bioethanol system requires organisms that together satisfy two complementary functions: efficient photosynthetic CO_2_ fixation and high-yield fermentative ethanol production. Microalgae, particularly members of the genus Chlorella, have emerged as the organisms of choice for the photosynthetic stage. Chlorella vulgaris is a unicellular green microalga that grows photoautotrophically using light energy, CO_2_, water, and inorganic nutrients to produce biomass rich in carbohydrates, lipids, and proteins [1,3].

The biological performance of C. vulgaris is characterized by CO_2_ fixation rates of 1.5–2.5 g CO_2_ per g dry biomass per day, a carbohydrate content that can reach 50% of dry cell weight under nitrogen-limited conditions, and a lipid content of 20–60% of dry weight depending on growth conditions [1,4]. These properties make C. vulgaris suitable for both carbohydrate extraction for bioethanol and lipid extraction for biodiesel production. In the present platform, the carbohydrate fraction is the primary product of interest, as it serves as the substrate for downstream fermentation.

C. vulgaris growth is governed by photosynthesis, carbon-concentrating mechanisms (CCMs), and lipid biosynthesis pathways. Under normal CO_2_ availability, RuBisCO in the Calvin cycle fixes CO_2_ into three-carbon compounds that are subsequently condensed into glucose and other carbohydrates. Under nitrogen or phosphorus limitation, carbon flux is redirected from protein synthesis toward carbohydrate and lipid storage, a metabolic shift that can be exploited to enhance substrate accumulation for fermentation [3].

For the fermentation stage, Saccharomyces cerevisiae was selected as the fermenting microorganism. S. cerevisiae is the dominant industrial ethanol producer, capable of tolerating ethanol concentrations up to 15% (v/v), operating under anaerobic conditions, and achieving ethanol yields approaching the theoretical maximum of 0.51 g ethanol per g glucose (Gay-Lussac yield) [4,9]. The fermentation reaction proceeds according to the stoichiometry: C_6_H_12_O_6_ → 2 C_2_H_5_OH + 2 CO_2_, releasing CO_2_ that is captured and recycled to the bioreactor.

Together, C. vulgaris and S. cerevisiae form a complementary biological consortium: the algae generate the carbohydrate substrate while simultaneously capturing CO_2_ from industrial sources, and the yeast converts that substrate to bioethanol while releasing CO_2_ that is fed back to the algae, creating a partially closed carbon loop. This integrated design is central to the bioprocess modeled by the simulation platform.

### 2.2. System Design and Process Flow

The overall system comprises four principal subsystems: (1) the Photochemical Bioreactor, in which C. vulgaris converts CO_2_ to glucose; (2) the Fermenter, in which S. cerevisiae converts glucose to bioethanol; (3) a Wastewater Recycling System for sustainable water management; and (4) a CO_2_ Recycling System that captures fermentation off-gas and reintroduces it into the bioreactor.

#### 2.2.1. Photochemical Bioreactor

An airlift bioreactor constructed from acrylic glass was selected for the photosynthetic cultivation stage. This design was chosen for its superior gas–liquid mass transfer characteristics relative to stirred-tank reactors and its transparent walls that allow uniform light penetration. The working volume ranges from 5–100 L at laboratory scale and 500–5,000 L at industrial scale. A full-spectrum LED light source operating at 400–700 nm wavelength with adjustable intensity (100–300 μmol photons/m^2^/s) supports photosynthetic activity. A 16:8 light/dark photoperiod is imposed to simulate natural diel cycles and enhance cellular productivity. Environmental conditions are precisely controlled: temperature is maintained at 25–30°C via a water-jacket heating/cooling system, and pH is regulated at 6.5–7.5 through automated acid/base dosing. CO_2_-enriched gas (sourced from industrial flue gas) is sparged into the reactor at near-atmospheric pressure, with CO_2_ concentrations of 2–15% (v/v) to enhance carbon availability while avoiding excessive acidification.

#### 2.2.2. Biomass Processing and Hydrolysis

Following cultivation, the harvested microalgal biomass undergoes a processing step prior to fermentation. Biomass is first concentrated through sedimentation or centrifugation and subsequently subjected to cell disruption and hydrolysis to release fermentable sugars from intracellular carbohydrates. Acid or enzymatic hydrolysis converts stored polysaccharides into glucose, which serves as the substrate for downstream fermentation by Saccharomyces cerevisiae. In the present simulation platform, this step is simplified by assuming a fixed carbohydrate-to-glucose conversion efficiency.

#### 2.2.3. Fermenter

The fermentation stage employs a stainless-steel stirred-tank bioreactor with a working volume of 5–50 L (laboratory) or 500–5,000 L (industrial). The tank utilizes 70–80% of its total volume to accommodate foaming and provide headspace. Two Rushton turbine impellers and four vertical baffles ensure uniform mixing, with agitation at 100–300 rpm. Temperature is maintained at 30–35°C via a thermal jacket, and pH is controlled at 4.5–5.5 using automated dosing. Anaerobic conditions are sustained by sparging CO_2_ from the recycling system, preventing oxygen intrusion. Fermentation duration is typically 24–72 h, depending on the desired ethanol concentration.

#### 2.2.4. CO_2_ Recycling Subsystem

Fermentation off-gas, consisting primarily of CO_2_ with trace ethanol vapors, is directed to a dedicated capture unit. A cooling condenser separates ethanol from the gas stream, followed by a scrubber that removes residual volatile organic compounds. The purified CO_2_ is then compressed to 1–2 bar using a compressor and stored in a buffer tank before controlled injection into the bioreactor via a flow-control valve. A feedback loop using in-line pH sensors adjusts CO_2_ injection rates dynamically to maintain target bioreactor pH. This closed-loop design reduces the net CO_2_ footprint of the process and lowers fresh CO_2_ demand.

**Figure 2.**
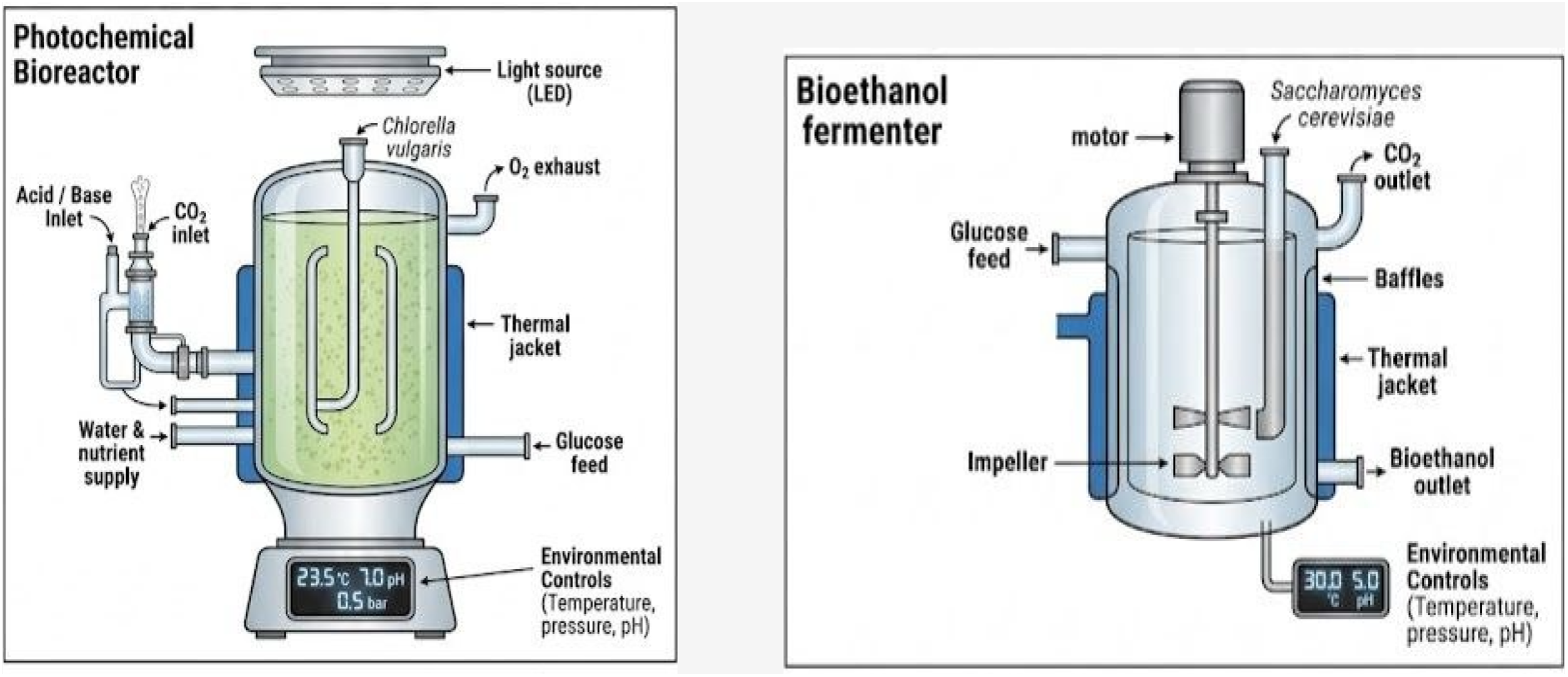
Engineering illustrations of the photochemical bioreactor (left) and bioethanol fermenter (right), showing key components including CO_2_ inlet, LED light source, thermal jacket, impellers, and environmental controls.

### 2.3. Bioprocess Mathematical Modeling

The simulation engine integrates three established kinetic models that collectively describe biomass growth, substrate consumption, and product formation across both biological stages of the process. In the present model, Monod kinetics describe substrate limitation, while the logistic growth term accounts for environmental carrying capacity, resulting in a combined formulation that captures both nutrient limitation and density-dependent growth effects.

#### 2.3.1. Monod Model for Substrate-Limited Growth

The Monod model, originally derived from observations of bacterial growth kinetics [5], relates the specific growth rate (μ) to the concentration of a single limiting substrate (S):

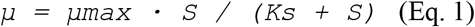

where μmax is the maximum specific growth rate (h^⁻1^), S is the substrate concentration (g/L), and Ks is the half-saturation constant (g/L), representing the substrate concentration at which μ = μmax/2. In the photosynthetic bioreactor, CO_2_ is the limiting substrate; in the fermenter, glucose (derived from algal carbohydrates) is the limiting substrate. The Monod model was applied to both Chlorella vulgaris and Saccharomyces cerevisiae growth with organism-specific parameter values (Table 2).

**Table 1.**
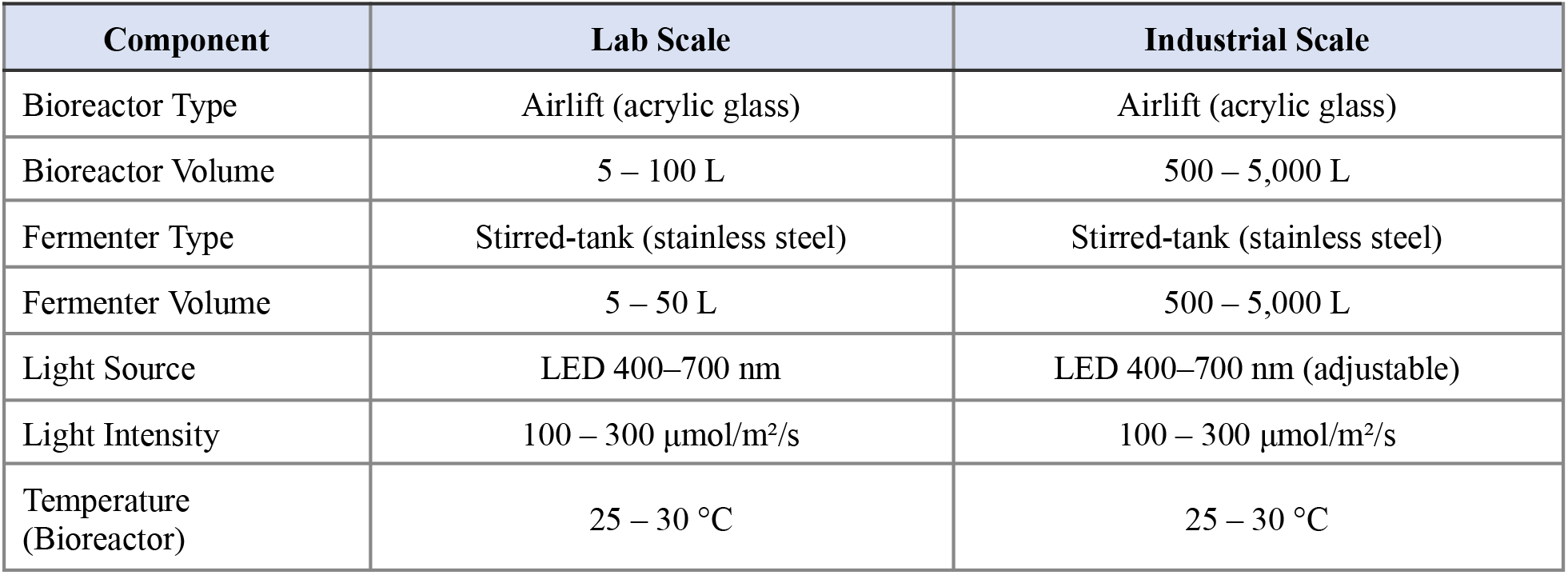
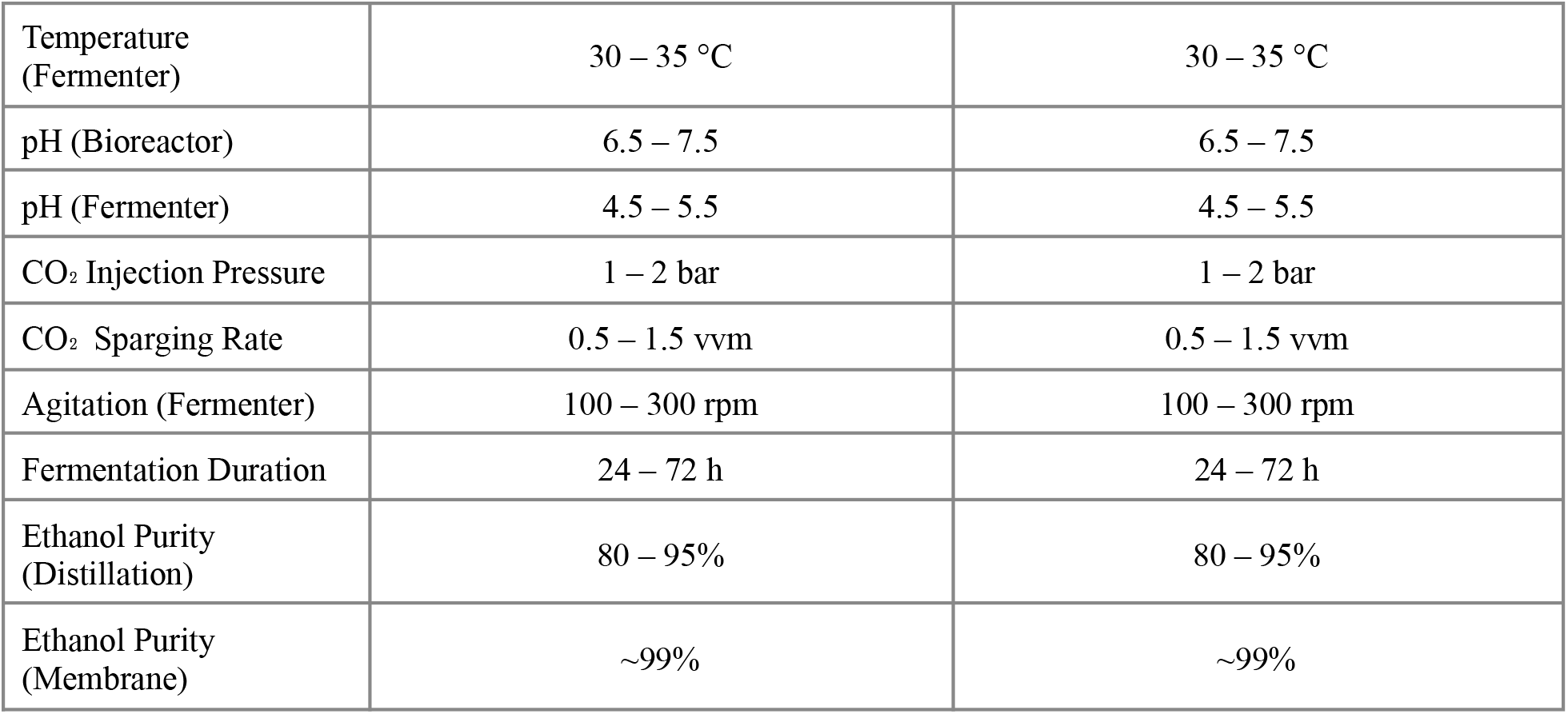
Design specifications of the bioreactor and fermenter at laboratory and industrial scales.

| Component | Lab Scale | Industrial Scale |
| --- | --- | --- |
| Bioreactor Type | Airlift (acrylic glass) | Airlift (acrylic glass) |
| Bioreactor Volume | 5 – 100 L | 500 – 5,000 L |
| Fermenter Type | Stirred-tank (stainless steel) | Stirred-tank (stainless steel) |
| Fermenter Volume | 5 – 50 L | 500 – 5,000 L |
| Light Source | LED 400–700 nm | LED 400–700 nm (adjustable) |
| Light Intensity | 100 – 300 $\mu\text{mol}/\text{m}^2/\text{s}$ | 100 – 300 $\mu\text{mol}/\text{m}^2/\text{s}$ |
| Temperature (Bioreactor) | 25 – 30 °C | 25 – 30 °C |
| Temperature (Fermenter) | 30 – 35 °C | 30 – 35 °C |
| pH (Bioreactor) | 6.5 – 7.5 | 6.5 – 7.5 |
| pH (Fermenter) | 4.5 – 5.5 | 4.5 – 5.5 |
| CO<sub>2</sub> Injection Pressure | 1 – 2 bar | 1 – 2 bar |
| CO<sub>2</sub> Sparging Rate | 0.5 – 1.5 vvm | 0.5 – 1.5 vvm |
| Agitation (Fermenter) | 100 – 300 rpm | 100 – 300 rpm |
| Fermentation Duration | 24 – 72 h | 24 – 72 h |
| Ethanol Purity (Distillation) | 80 – 95% | 80 – 95% |
| Ethanol Purity (Membrane) | ~99% | ~99% |

**Table 2.** Kinetic model parameters used in the simulation engine, with literature-based ranges.

| Parameter | Symbol | Value / Range | Units |
| --- | --- | --- | --- |
| Max. algal growth rate | $\mu_{max,algae}$ | 0.1 – 0.3 | $h^{-1}$ |
| Max. yeast growth rate | $\mu_{max,yeast}$ | 0.05 – 0.5 | $h^{-1}$ |
| Half-saturation constant | Ks | 0.001 – 0.1 | g/L |
| Max. algal biomass | $X_{max,algae}$ | 5 – 20 | g/L |
| Max. yeast biomass | $X_{max,yeast}$ | 10 – 50 | g/L |
| Growth-assoc. coefficient | $\alpha$ | 0.1 – 0.5 | g/g |
| Non-growth-assoc. coeff. | $\beta$ | 0.01 – 0.1 | g/(g·h) |
| CO<sub>2</sub> yield coefficient | $Y_{CO_2/X}$ | 1.8 - 2.2 | g CO<sub>2</sub>/g biomass |
| Biomass yield | $Y_{biomass}$ | 0.3 – 0.6 | g/g |
| Ethanol yield | $Y_{ethanol}$ | 0.4 – 0.51 | g/g |

#### 2.3.2. Logistic Growth Model for Carrying Capacity

To capture density-dependent growth limitations, the logistic model [6] was incorporated as a carrying-capacity constraint applied to Monod growth kinetics.

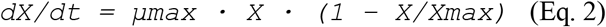

where X is the biomass concentration (g/L), Xmax is the carrying capacity (maximum biomass concentration supportable by the culture environment, g/L), and t is time (h). This equation yields a sigmoidal growth curve that transitions from near-exponential growth at low biomass densities to a stationary phase as X approaches Xmax. The analytical solution of Equation 2 is:

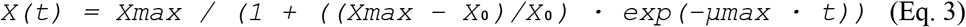

where X_0_ is the initial biomass concentration at t = 0. This formulation captures the four characteristic phases of microbial batch culture: lag (not explicitly modeled but reflected in X_0_ initialization), exponential and stationary (represented by the asymptotic approach to Xmax). Characteristic parameter ranges are Xmax = 5–20 g/L for C. vulgaris and 10–50 g/L for S. cerevisiae.

#### 2.3.3. Luedeking-Piret Model for Product Formation

Ethanol production by S. cerevisiae is modeled using the Luedeking-Piret framework [7,8], which partitions product formation into a growth-associated component (proportional to the rate of biomass increase) and a non-growth-associated component (proportional to current biomass concentration):

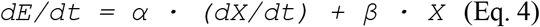

where E is the ethanol concentration (g/L), α is the growth-associated product formation coefficient (g ethanol/g biomass), and β is the non-growth-associated coefficient (g ethanol/(g biomass · h)). The parameter α captures the metabolic coupling between growth and ethanol synthesis, while β accounts for maintenance metabolism-driven ethanol production independent of active growth. For C. vulgaris, carbohydrate accumulation was approximated using a simplified production model proportional to biomass growth, representing intracellular carbohydrate storage rather than extracellular secretion.

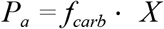

Where: *f*_*carb*_ = carbohydrate fraction of biomass

#### 2.3.4. CO_2_ Mass Balance

CO_2_ consumption in the bioreactor is linked to algal biomass production through a yield coefficient:

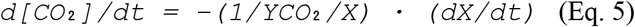

where YCO_2_/X is the CO_2_ yield coefficient (g CO_2_ consumed per g biomass produced). This relationship enables real-time tracking of carbon utilization efficiency throughout the simulation.

To maintain physical realism, the simulation tracks carbon flows between CO_2_, biomass, glucose, and ethanol pools. Carbon conservation is enforced through yield coefficients linking biomass formation, carbohydrate accumulation, and ethanol production.

#### 2.3.5. Efficiency Metrics

Three efficiency metrics are computed by the platform to provide a multi-dimensional assessment of system performance:

Molar efficiency (η_mol) measures the mole-to-mole conversion of net CO_2_ consumed into ethanol:

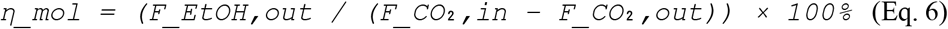

Mass yield efficiency (η_mass) expresses grams of ethanol produced per gram of CO_2_ fed to the system:

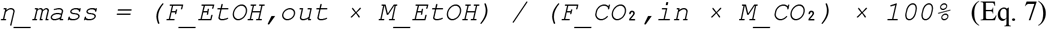

Overall process efficiency (η_total) represents the percentage of the theoretical maximum conversion yield achieved by the modeled system where the theoretical maximum is 0.348 g ethanol/g CO_2_.

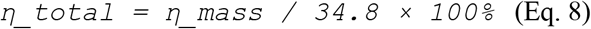

where M_EtOH = 46 g/mol and M_CO_2_ = 44 g/mol. These formulations follow established bioprocess engineering conventions [14,15].

### 2.4. Integration of the Two-Phase Model

The two biological phases are coupled through substrate linkage: carbohydrates produced by C. vulgaris (Pa, g/L) in the photosynthetic bioreactor serve directly as the glucose substrate (S) for S. cerevisiae in the fermenter. The simulation time-steps through each phase sequentially, transferring the output carbohydrate concentration from Phase 1 as the input substrate concentration for Phase 2. Key variable attributes tracked throughout include biomass concentration (algal and yeast), substrate concentration (CO_2_ and glucose), product concentration (ethanol), CO_2_ evolution rate, and process efficiency.

Environmental control algorithms are embedded as correction factors applied to the base kinetic rates: temperature effects follow Arrhenius-based activation energy dynamics; pH effects are represented by sigmoidal response functions; light availability in the bioreactor is modeled using the Lambert-Beer law for light attenuation through the culture; and nutrient limitation is incorporated through Liebig’s law of the minimum, ensuring that the most limiting factor governs the overall growth rate.

## 3. Software Architecture and Technical Implementation

### 3.1. System Requirements and Design Philosophy

The platform was designed as a portable, standalone desktop application to maximize accessibility for academic and research users who may not have access to commercial simulation suites. The application runs without requiring a web browser, operates across Windows, macOS, and Linux, and executes the simulation backend in under 5 seconds on standard hardware (4–8 GB RAM, any modern CPU), compared to the ~2-minute execution time observed when the original MATLAB implementation was used.

The development followed an Incremental Agile methodology, with two software developers (A. Mirghani and T. Omer) working in parallel on frontend and backend components respectively. This approach allowed iterative testing and integration of each simulation module before incorporation of the next, reducing integration errors and enabling continuous validation of mathematical outputs against expected bioprocess behavior.

### 3.2. Frontend Architecture

The user interface was built entirely in JavaScript using React (v19.1.0) for component-based UI rendering and React Router DOM (v7.5.2) for client-side navigation. The application is wrapped in Electron (v35.1.5) to compile the web-based interface into a native desktop executable, with Vite (v6.2.0) providing fast bundling and hot-module replacement during development. Styling combines utility-first Tailwind CSS classes with custom CSS for fine-grained layout control.

Three-dimensional visualization of the bioreactor and fermenter models is implemented using Three.js (v0.176.0) in combination with @react-three/fiber (v9.1.2) as a React renderer and @react-three/drei (v10.0.7) for abstraction helpers. Framer Motion (v12.12.1) handles UI animations, including transitions between pages and loading indicators. Lucide React and React Icons provide the iconography. The application comprises five principal views: (1) Homepage, (2) Input Page, (3) Output Page, (4) 3D Models Page, and (5) Simulation History.

**Figure 3.**
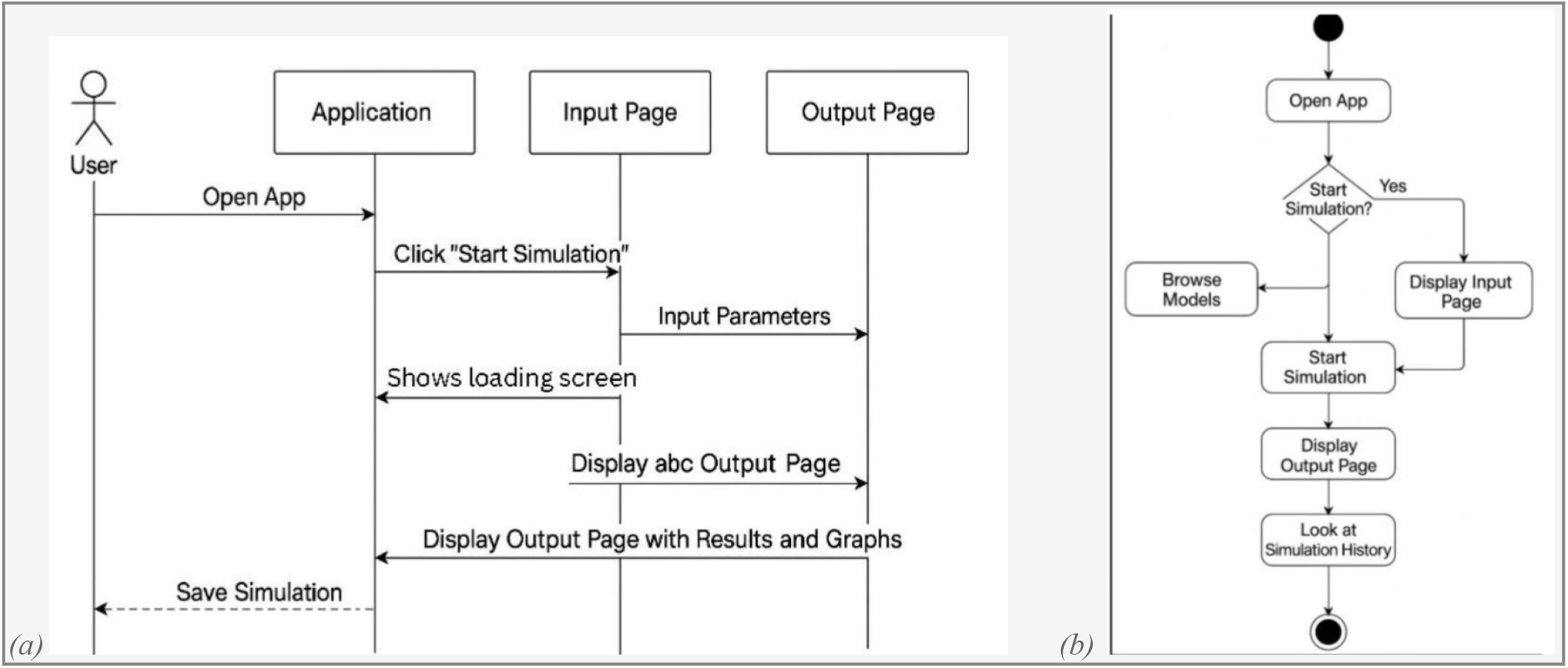
(a) User interaction sequence diagram showing the flow from application launch to simulation output and history storage. (b) State-transition flowchart illustrating possible user pathways through the application.

### 3.3. Backend Architecture

The simulation engine was initially prototyped in MATLAB 2018, which provided a convenient environment for equation development and validation. However, MATLAB requires installation of licensed software on each target machine, and the compiled MATLAB executable exhibited unacceptably long execution times (~2 minutes per simulation run). The engine was therefore re-implemented in Python 3.13, utilizing NumPy for numerical integration and Matplotlib for graph generation. The Python script executes in approximately 4 seconds per simulation run—a 30-fold reduction compared to MATLAB—and was compiled into a Windows-compatible executable using PyInstaller.

Electron’s Inter-Process Communication (IPC) mechanism manages communication between the JavaScript frontend and the Python executable backend. When the user submits simulation parameters via the Input Page, Electron’s main process invokes the Python executable via execFile(), passing parameters as command-line arguments. The Python backend writes simulation results (numerical summary as a text file), graphs (as PNG images), and a formatted PDF report to a designated output folder in the application root directory. The React frontend then reads and renders these outputs on the Output Page. Simulation history is maintained as a collection of PDF reports accessible through the History module.

## 4. Results

### 4.1. Simulation Scenario Analysis

Three representative simulation runs were conducted to assess platform behavior across a range of operating conditions. The input parameters and corresponding outputs for each run are summarized in Table 3. The runs were deliberately selected to span contrasting regimes: a moderate short-duration run (Run 1), a high-biomass extended run (Run 2), and a minimal short-duration run (Run 3).

**Table 3.** Input parameters and simulation output metrics for three representative simulation scenarios.

| Output Metric | Run 1 | Run 2 | Run 3 | Unit |
| --- | --- | --- | --- | --- |
| Initial Algal Biomass | 0.30 | 0.50 | 0.30 | g/L |
| Initial Yeast Biomass | 0.50 | 15.0 | 0.10 | g/L |
| Initial CO<sub>2</sub> | 0.05 | 0.50 | 0.05 | g/L |
| Simulation Duration | 5.00 | 24.0 | 1.00 | h |
| Total Glucose Produced | 2.38 | 14.75 | 0.04 | g/L |
| Final Algal Biomass | 5.03 | 30.00 | 0.18 | g/L |
| Total Ethanol Produced | 0.75 | 6.64 | 0.00 | g/L |
| Total CO<sub>2</sub> Produced | 0.80 | 7.14 | 0.05 | g/L |
| Process Efficiency | 93.78 | 92.99 | 1.09 | % |

Run 1 (5 hours, moderate inocula) demonstrated that the platform correctly resolves the exponential-to-stationary growth transition of C. vulgaris, with algal biomass increasing from 0.30 g/L to 5.03 g/L—approaching the logistic model’s carrying capacity. Total glucose production of 2.38 g/L provided sufficient substrate for yeast fermentation, yielding 0.75 g/L ethanol. The high process efficiency of 93.78% relative to the theoretical maximum conversion efficiency in this run reflects near-complete substrate utilization under the given conditions and validates the internal consistency of the coupled kinetic model framework.

Run 2 (24 hours, high yeast inoculum at 15 g/L) represents an industrial-scale scenario with prolonged fermentation. Algal biomass reached 30.00 g/L, generating 14.75 g/L glucose and 6.64 g/L ethanol—values consistent with reported experimental ranges for S. cerevisiae fermentation of algal hydrolysates [2,5]. The slight reduction in efficiency to 92.99% compared to Run 1 is attributed to increased CO_2_ production (7.14 g/L) from the larger yeast biomass, which raises the denominator in the efficiency calculation.

Run 3 (1 hour, minimal conditions) reveals the platform’s ability to resolve low-efficiency regimes: with insufficient biomass development and substrate accumulation, essentially no ethanol was produced (0.00 g/L) and process efficiency collapsed to 1.09%. This result correctly captures the behavior of a system that has not yet progressed beyond the lag phase, confirming that the system remains in an early growth regime where biomass accumulation is minimal due to short simulation duration.

### 4.2. Efficiency Analysis

Applying the three efficiency metrics to an illustrative scenario (FCO_2_,in = 120 mol, FCO_2_,out = 40 mol, FEtOH,out = 20 mol) demonstrates the complementary information provided by each formulation (Table 4). The molar efficiency of 25.0% indicates that one-quarter of the net CO_2_ consumed was converted to ethanol on a molar basis. The overall efficiency of 50% corresponds to 50% of the theoretical maximum yield of 34.8 g ethanol per 100 g CO_2_, suggesting that the modeled system operates in a feasible but not yet optimized regime. These metrics provide researchers with actionable targets for parameter optimization.

**Table 4.**
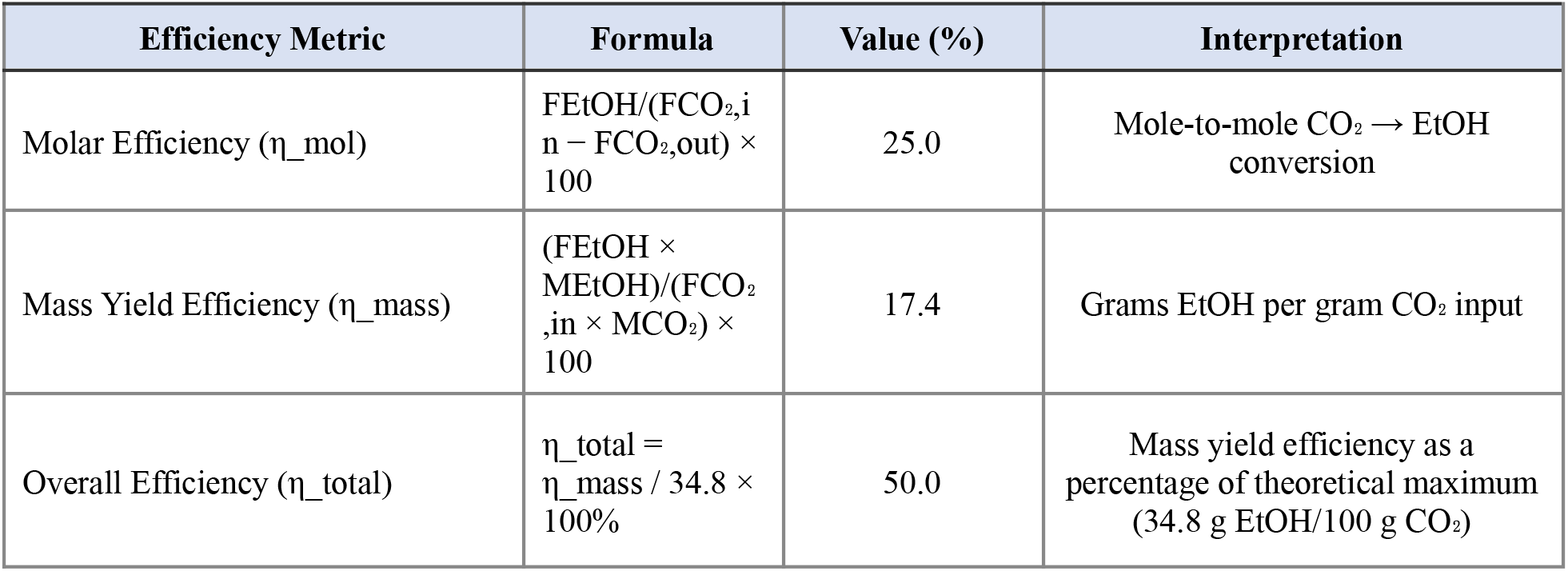
Efficiency metric calculations for an illustrative simulation scenario (F_CO_2_,in = 120 mol, F_CO_2_,out = 40 mol, F_EtOH,out = 20 mol).

| Efficiency Metric | Formula | Value (%) | Interpretation |
| --- | --- | --- | --- |
| Molar Efficiency ($\eta_{mol}$) | $\frac{F_{EtOH}}{(F_{CO_{2,i}} - F_{CO_{2,o}})} \times 100$ | 25.0 | Mole-to-mole $CO_2 \rightarrow EtOH$ conversion |
| Mass Yield Efficiency ($\eta_{mass}$) | $\frac{(F_{EtOH} \times M_{EtOH})}{(F_{CO_{2,i}} \times M_{CO_2})} \times 100$ | 17.4 | Grams EtOH per gram $CO_2$ input |
| Overall Efficiency ($\eta_{total}$) | $\eta_{total} = \eta_{mass} / 34.8 \times 100\%$ | 50.0 | Mass yield efficiency as a percentage of theoretical maximum (34.8 g EtOH/100 g $CO_2$) |

**Figure 4.**
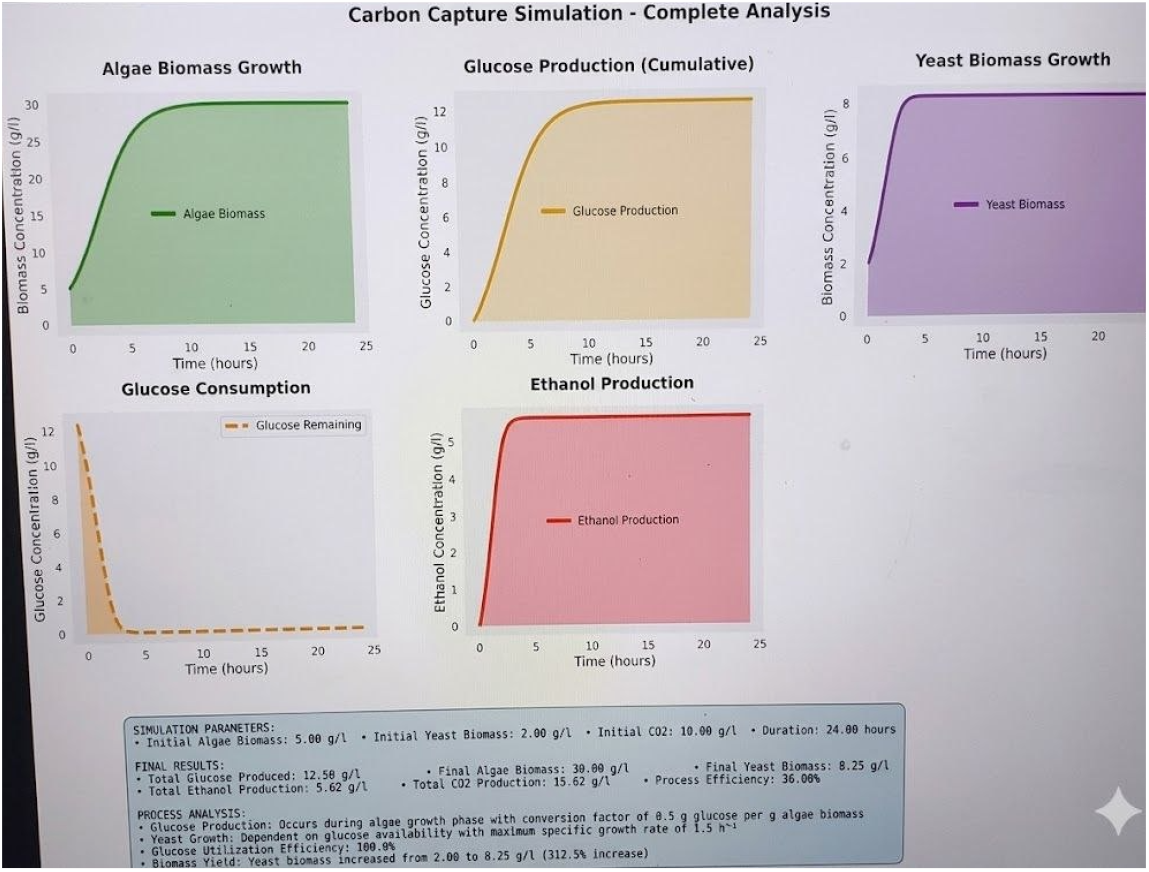
Representative simulation output curves: (a) algal biomass growth following logistic kinetics; (b) glucose production; (c) yeast biomass growth following logistic kinetics; (d) glucose consumption by yeast (e) ethanol production.

### 4.3. Platform Functionality and User Experience

The Output Page of the application renders five graphs (biomass vs. time, glucose/ethanol production, glucose consumption) alongside a textual summary of key output metrics. Users can navigate directly from the Output Page back to the Input Page to modify parameters and rerun the simulation. Each simulation run automatically generates a PDF summary report stored in the History module, allowing retrospective comparison of scenarios.

The Models Page provides interactive 3D renderings of the photochemical bioreactor and fermenter, allowing users to rotate, zoom, and inspect each component. This visualization feature is particularly valuable for educational contexts, enabling students to develop spatial understanding of bioprocess equipment before interacting with the mathematical simulation. The 3D models were created using Three.js and integrated into the React component tree via @react-three/fiber, with smooth rotation animations powered by Framer Motion.

## 5. Discussion

The Carbon Capture Modeling and Simulation Platform presented here represents a significant step toward accessible, integrated bioprocess simulation tools for carbon management research. By coupling the Monod, Logistic, and Luedeking-Piret models within a user-friendly desktop application, the platform enables rapid exploration of the parameter space governing CO_2_-to-bioethanol conversion without requiring expensive commercial licenses or deep programming expertise.

The simulation results across the three test scenarios demonstrate the platform’s sensitivity to initial conditions and process duration—key parameters that directly govern bioprocess performance in practice. The high efficiencies observed in Runs 1 and 2 (>92%) reflect near-complete substrate utilization under favorable conditions, while the near-zero efficiency of Run 3 correctly captures the behavior of a prematurely terminated or under-inoculated system. This range of behavior confirms that the mathematical framework embedded in the platform is physically consistent and behaviorally realistic within the validated parameter ranges.

Several technical constraints merit acknowledgement. The kinetic coefficients used in the simulation (μmax, Ks, α, β, YCO_2_/X) are derived from literature values for pure-culture systems operating under idealized conditions. In practice, co-culture dynamics, contamination, microbial evolution, and environmental noise introduce variability that deterministic models cannot capture. The platform does not currently implement stochastic modeling or Monte Carlo sensitivity analysis, meaning that uncertainty quantification—an important consideration for industrial pre-feasibility assessments—must be performed externally. Additionally, the simulation operates on a single-strain assumption for both organisms, which may not reflect the performance of mixed microbial communities or evolved strains adapted to industrial conditions.The current simulation focuses primarily on biological conversion efficiencies and does not explicitly account for energy consumption associated with LED lighting, gas compression, mixing, or downstream processing. Future versions of the platform will incorporate energy balance calculations to enable full techno-economic assessment of the system. A further limitation concerns the carbon accounting of the CO_2_ recycling subsystem. While the platform models a partially closed-loop system in which fermentation off-gas is recycled to the bioreactor, the current simulation does not explicitly quantify the fraction of total CO_2_ input that is sourced from recycling versus fresh industrial feed. As a result, the efficiency metrics reported in Table 3 and Table 4 reflect overall CO_2_-to-ethanol conversion but do not distinguish between recycled and externally supplied carbon. Future versions of the platform should incorporate a dedicated recycling efficiency metric - for example, the ratio of recycled CO_2_ to total CO_2_ input - to provide a more complete and transparent carbon balance. This addition would strengthen the platform’s utility for industrial pre-feasibility assessments where minimizing fresh CO_2_ demand is a primary design objective.

Despite these limitations, the platform occupies a valuable niche between overly simplified spreadsheet tools and full-scale commercial process simulators. For educational applications, it provides immediate, interactive feedback that bridges theoretical kinetics with system-level understanding. For pre-feasibility research, it enables rapid screening of operating conditions before committing to expensive experimental campaigns. Future enhancements should include stochastic variability modules, integration with real-time sensor data streams for digital twin applications, and expansion to additional biofuel production pathways (e.g., biodiesel from algal lipids, biogas from anaerobic digestion of fermentation residues).

The choice of an Electron/React/Python technology stack—while unconventional for scientific computing—proved highly effective for this application. The 30-fold reduction in simulation execution time achieved by migrating from MATLAB to Python, combined with the elimination of external software dependencies through PyInstaller compilation, significantly lowers the barrier to deployment and use. The cross-platform Electron wrapper ensures that the application can be distributed to Windows, macOS, and Linux users without modification, broadening the potential user base well beyond institutions with access to MATLAB licenses.

## 6. Conclusions

This work presented the design, mathematical formulation, and technical implementation of a Carbon Capture Modeling and Simulation Platform that models the full conversion pathway from industrial CO_2_ to bioethanol via coupled microalgal and yeast bioprocesses. The principal findings and contributions can be summarized as follows:

- The platform integrates three validated kinetic models (Monod, Logistic, Luedeking-Piret) within a closed-loop CO_2_ recycling framework, enabling system-level simulation of a CO_2_-to-bioethanol process from a single user interface.
- Three simulation scenarios spanning a wide range of input conditions yielded process efficiencies from 1.09% to 93.78%, confirming the platform’s sensitivity to initial conditions and its ability to resolve both favorable and unfavorable operating regimes.
- Migration of the simulation backend from MATLAB to Python (via PyInstaller) reduced execution time from ~2 minutes to ~4 seconds, while eliminating the requirement for licensed software on end-user machines.
- The Electron/React frontend provides interactive 3D bioreactor visualizations, real-time graphical outputs, and a simulation history module, making the tool suitable for both research and educational contexts.
- The efficiency analysis demonstrates that modeled systems can approach 50% of the theoretical maximum CO_2_-to-ethanol conversion yield, establishing a quantitative benchmark for further process optimization.

The platform represents a foundational tool for the pre-feasibility assessment of integrated carbon capture and biofuel production systems. Future work will focus on incorporating stochastic modeling for uncertainty quantification, extending the biological scope to include lipid-based biodiesel pathways and anaerobic digestion, and developing a web-based deployment option to further broaden accessibility. The authors anticipate that continued development will yield a comprehensive simulation suite capable of supporting both academic research and early-stage industrial process design in the emerging field of biological carbon capture and utilization.

## Acknowledgments

The authors express sincere gratitude to their supervisor, Asst. Prof. Dr. Nihal Bayir (Department of Bioengineering, Cyprus International University), for her guidance, constructive feedback, and support throughout the development of this project. The authors also thank the Faculty of Engineering at Cyprus International University for providing the academic environment and resources that facilitated this work.

## Funding

There are no sources of external funding to declare for this project.

## Author Contributions

Conceptualization and bioprocess design: A. Hamid; Literature review and organism selection: N. Akasha; Bioprocess modeling (mass and energy balances): P. Kapenda Mukumbi; Simulation engine development and software architecture: A. Mirghani and T. Omer; Supervision: Dr. N. Bayir. All authors reviewed and approved the final manuscript.

## Conflict of Interest

The authors declare no conflicts of interest.

## Data Availability Statement

Simulation data and platform source code are available upon request from the corresponding author. The compiled application executable is available for distribution to academic users.

## Notes

### Competing Interest Statement

The authors have declared no competing interest.

